# Yin Yang Interactomes Drive the Metabolic Dysfunction-Associated Liver Disease Continuum to Hepatocellular Carcinoma

**DOI:** 10.64898/2026.01.06.698028

**Authors:** Tyler L. Bissoondial, Ravi Reddi, Sita Sirisha Madugula, Manisha Shukla, Mahesh Narayan, Prakash Narayan

## Abstract

The risk for hepatocellular carcinoma (HCC) arising from metabolic dysfunction– associated steatotic liver disease (MASLD)–related cirrhosis exceeds the risk for cancer developing in the kidney, heart, or lung as a consequence of end-stage disease in those organs. Both experimental and clinical studies support the existence of circulating biomarkers whose elevated expression is associated with, and supports the diagnosis of, HCC. We posit the existence of oncogenic HCC biomarkers under regulation by tumor-suppressive microRNAs (miRs), and that reduced tumor-suppressive miR expression leads to increased oncogenic biomarker expression. By interrogating the MASLD–HCC miR interactome and miRs known to regulate HCC biomarkers, we identified hsa-miR-577, hsa-miR-9500, hsa-miR-101-3p, miR-206, hsa-miR-219a-1-3p, and hsa-miR-613 as tumor-suppressive miRs whose expression is reduced across multiple cancers. Their target genes, osteopontin, Golgi protein-73, and Dickkopf-related protein 1, exhibit increased expression in HCC and display oncogenic activity. Collectively, dysregulated expression of these miRs and their downstream gene products may contribute to progression along the MASLD continuum toward HCC.

**Highlights:** We identified tumor suppressive miRs, hsa-miR-577, hsa-miR-9500, hsa-miR-101-3p, miR-206, hsa-miR-219a-1-3p, and hsa-miR-613 with potentially reduced expression levels in HCC. Their gene products, osteopontin, Golgi protein-73, and Dickkopf-related protein 1, exhibit oncogenic properties and increased expression levels in HCC. These miR-gene product interactions may drive the MASLD continuum to HCC. Mechanistic insights should illuminate therapies. Identification of therapeutics that interrupt these interactions may represent a therapeutic pillar for patients with MASLD.

Figure
MASLD-related HCC.
The intersection set between the MASLD-HCC miR interactome and the set of miRs regulating biomarkers diagnostic for HCC was interrogated and tumor suppressive miRs regulating HCC biomarkers with oncogenic activity were identified. Increased risk for HCC in patients with MASLD and MASLD-cirrhosis may arise from interactions between these dysregulated miRs and their downstream gene products.

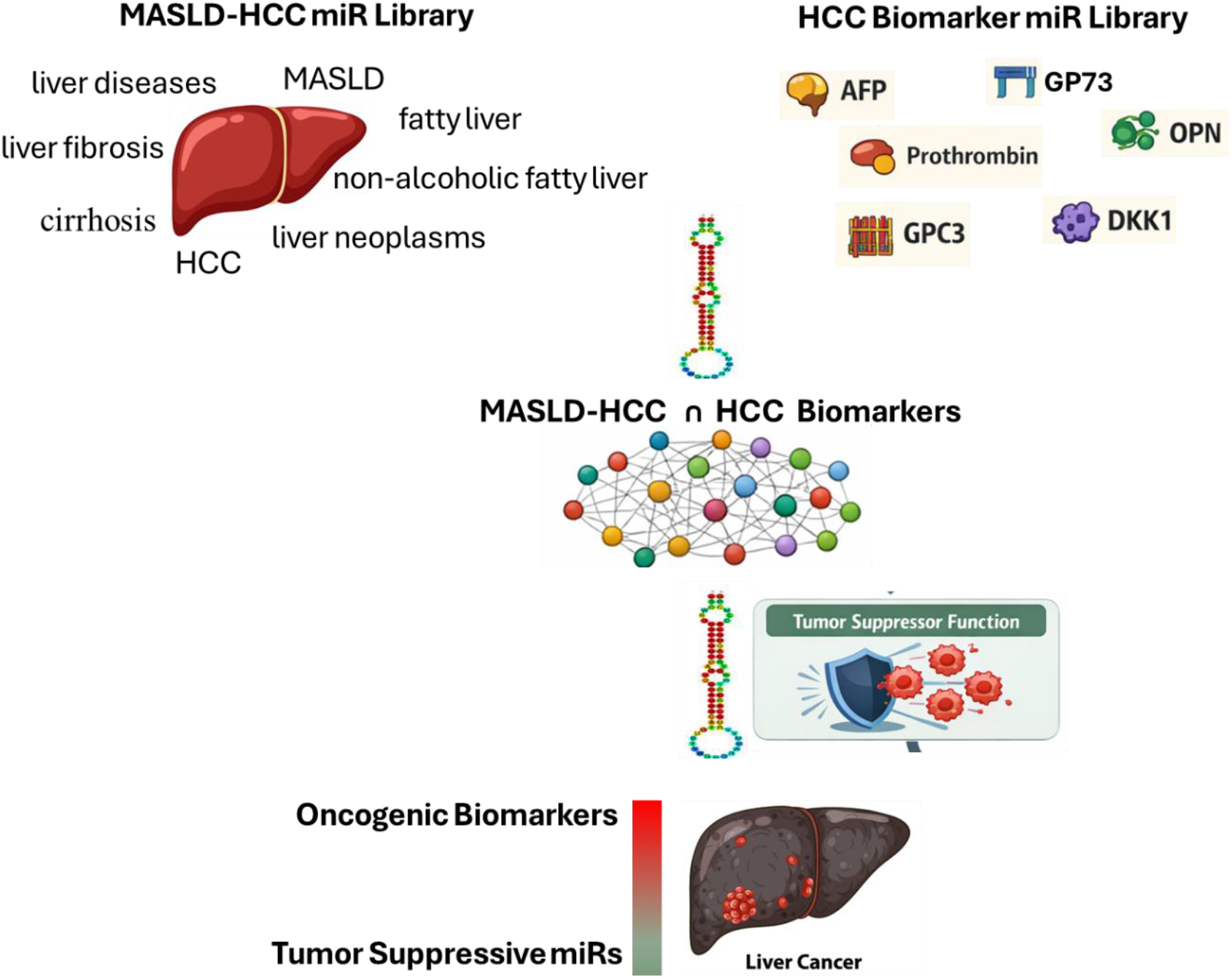

**Prologue:** Metabolic Dysfunction-Associated Liver Disease (MASLD), the leading cause of hepatocellular carcinoma (HCC) in Sweden, is now taking epidemic proportions in South Asia. In India, there is a relatively high prevalence of pre-diabetes, diabetes and insulin resistance; body mass indices > 35 are being observed. Each of these and/or their combinations represent risk factors for MASLD. Nevertheless, both patients and providers are largely oblivious to this disease, both from a lack of general awareness and the asymptomatic nature of MASLD, at least early within its continuum. Diagnosis is often made late, and by exclusion, when presentation may include fatigue, drowsiness, mild encephalopathy (MASLD-pituitary feedback loop), prominent hepatomegaly and asciites, and with liver-specific test demonstrating stiffness, scarring and even HCC. Societal awareness and effective management of this disease are urgent needs. A strategy akin to the 4 pillars for treatment of type 2 diabetes and chronic kidney disease (CKD) needs to be implemented for treatment of patients with MASLD.

Figure
Four-pillar Strategy for MASLD.
A strategy akin to the one in place for treatment of type 2 diabetes and CKD needs to be implemented for treatment of patients with MASLD. Aspects of this strategy, viz. SGLT2i and GLP-1±GIP agonists, are already approved in MASLD.

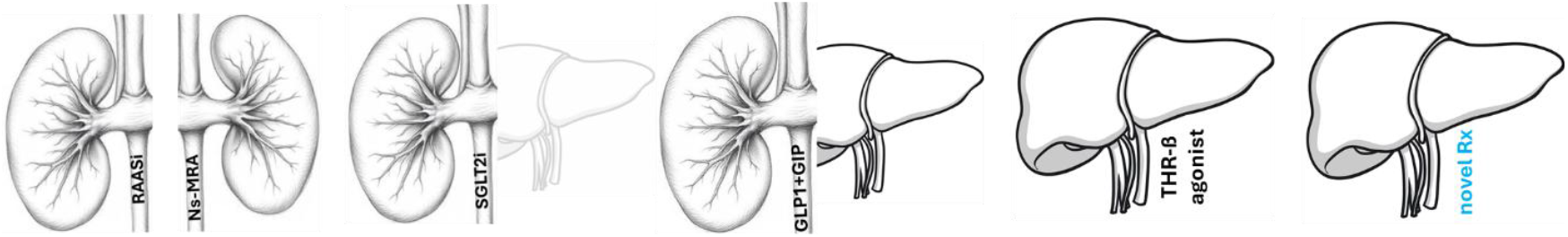

## Introduction

Metabolic dysfunction-associated steatotic liver disease (MASLD) affects ∼ 100 million adults in the US alone and ∼1.3 billion adults worldwide^1-4^. Driven in large part by the diabetes, obesity and metabolic syndrome epidemics^3,4^, MASLD starts with simple steatosis or accumulation of lipid droplets within the liver. When inflammation follows, the condition is termed metabolic dysfunction-associated steatohepatitis (MASH), a stage accompanied by an increase in liver enzymes^5^. The continuum can progress to MASH with fibrosis, and then cirrhosis, which is severe scarring of the liver, disfigurement of the organ and potential for decline in liver function^5^. Cirrhosis not only represents a risk for liver failure but also poses a significant risk for progression to hepatocellular carcinoma (HCC)^6^. Indeed, 80-90% of HCC develops in cirrhotic livers, albeit not all related to MASLD-cirrhosis^7^. Clinical data indicate that 20-30% of MASLD-related HCC cases occur in the absence of cirrhosis^8-12^. Given the sheer size of the metabolic disease epidemic, MASLD is projected to be the leading cause of HCC within the next few decades^13^.

Failed and/or maladaptive tissue repair and an epithelial-to-mesenchymal transition (EMT) represent fibrotic disease-associated mechanisms shared across multiple organs including the liver, kidney, lung and heart^14^. Interstitial lung disease and idiopathic pulmonary fibrosis affect hundreds of thousands of individuals in the United States alone and are major causes of morbidity and mortality, particularly among older adults^15^. The prevalence of chronic kidney disease is comparable to that of MASLD, and progressive renal scarring can culminate in end-stage renal disease, which affects millions globally^16^. Myocardial fibrosis is a defining feature of both idiopathic dilated cardiomyopathy and ischemic cardiomyopathy^17,18^. Unlike the MASLD-MASH-fibrosis-cirrhosis-HCC continuum, fibrosis of the lung, kidney, and heart do not carry a comparable increased risk of malignancy in the affected tissues. These statistics suggest that factors within the hepatic microenvironment drive MASLD towards HCC.

Both experimental^5^ and clinical findings^19^ inform multiple circulating biomarkers that support a diagnosis of HCC. These biomarkers include alpha fetoprotein (AFP), des γ carboxyprothrombin (DCP/PIVKA II), glypican-3, osteopontin (OPN), golgi protein-73 (GP73) and Dickkopf-related protein 1 (DKK1)^5^. Levels of these biomarkers are increased in HCC^5^, driven by increased expression of their mRNA which in turn is driven by a reduction in expression levels of microRNA (miRs) regulating these mRNA^20^. Our overarching hypothesis is there exist oncogenic HCC biomarkers under regulation by tumor-suppressive microRNAs (miRs), and that reduced tumor-suppressive miR expression leads to increased oncogenic biomarker expression. As a first step towards addressing this hypothesis we seek to identify tumor-suppressing specific miRs and their gene products.

## Methods

A MASLD–related HCC interactome was constructed using miRNet (miRNet) by querying the miR–disease module with the in-built terms liver neoplasms, liver fibrosis, liver diseases, cirrhosis, non-alcoholic fatty liver, non-alcoholic fatty liver disease, fatty liver, hepatocellular carcinoma HCC, and MASLD. This analysis generated a library of miRs associated with the MASLD–HCC continuum. Independently, the library of miRs predicted to regulate HCC biomarker genes, *afp* (AFP), *f2* (prothrombin), *spp1* (OPN), gpc3 (glypican-3), *golm1* (GP73), and *dkk1* (DKK1) was generated using miRDB (miRDB - MicroRNA Target Prediction Database). Those miRs that resided within the intersection of these two libraries were examined for tumor-suppressive activity based on the literature. Biomarker gene products associated with these tumor suppressive miRs were identified and examined for oncogenic potential.

## Results

The MASLD-related HCC miR interactome (Figure 1) encompasses different stages of the MASLD continuum, including fatty liver, fibrosis, cirrhosis and cancer, and miRs associated with these stages. More than 400 miRs comprise this interactome (Table 1).

**Table 1.** Library of miRs Associated with MASLD-related HCC.

|  |  |  |  |  |  |  |  |  |
| --- | --- | --- | --- | --- | --- | --- | --- | --- |
| hsa-let-7a-1 | hsa-mir-129-1 | hsa-mir-133a-2 | hsa-mir-363 | hsa-mir-193b | hsa-miR-33a-5p | hsa-mir-942 | hsa-mir-498 | hsa-mir-5 |
| hsa-let-7a-2 | hsa-mir-148a | hsa-mir-135a-1 | hsa-mir-365a | hsa-mir-497 | hsa-miR-92a-3p | hsa-mir-506 | hsa-mir-4488 | hsa-mir-6499 |
| hsa-let-7a-3 | hsa-mir-30d | hsa-mir-135a-2 | hsa-mir-302b | hsa-mir-181d | hsa-miR-101-3p | hsa-mir-129 | hsa-mir-3677 | hsa-mir-4715 |
| hsa-let-7b | hsa-mir-139 | hsa-mir-140 | hsa-mir-376c | hsa-mir-525 | hsa-miR-29b-3p | hsa-mir-320d-1 | hsa-mir-4746 | hsa-mir-6766 |
| hsa-let-7c | hsa-mir-10a | hsa-mir-141 | hsa-mir-369 | hsa-mir-524 | hsa-miR-103a-3p | hsa-mir-34 | hsa-mir-181a | hsa-mir-467b |
| hsa-let-7d | hsa-mir-10b | hsa-mir-142 | hsa-mir-370 | hsa-mir-519a-1 | hsa-miR-106a-5p | hsa-mir-1266 | hsa-mir-218 | hsa-mir-40 |
| hsa-let-7e | hsa-mir-34a | hsa-mir-143 | hsa-mir-372 | hsa-mir-500a | hsa-mir-107 | hsa-mir-199 | hsa-mir-1 | hsa-mir-44 |
| hsa-let-7f-1 | hsa-mir-181a-2 | hsa-mir-144 | hsa-mir-373 | hsa-mir-503 | hsa-miR-198 | hsa-mir-6838 | hsa-mir-125b | hsa-mir-466g |
| hsa-let-7f-2 | hsa-mir-181b-1 | hsa-mir-145 | hsa-mir-374a | hsa-mir-513a-1 | hsa-miR-199a-5p | hsa-mir-124a | hsa-mir-194 |  |
| hsa-mir-15a | hsa-mir-181c | hsa-mir-152 | hsa-mir-377 | hsa-mir-513a-2 | hsa-miR-199a-3p | hsa-mir-130 | hsa-mir-138 |  |
| hsa-mir-16-1 | hsa-mir-182 | hsa-mir-191 | hsa-mir-378a | hsa-mir-532 | hsa-mir-182 | hsa-mir-196 | hsa-mir-16 |  |
| hsa-mir-17 | hsa-mir-183 | hsa-mir-9-1 | hsa-mir-379 | hsa-mir-455 | hsa-mir-137 | hsa-mir-209 | hsa-mir-92a |  |
| hsa-mir-18a | hsa-mir-187 | hsa-mir-9-2 | hsa-mir-380 | hsa-mir-92b | hsa-mir-9 | hsa-mir-922 | hsa-mir-29b |  |
| hsa-mir-19a | hsa-mir-196a-2 | hsa-mir-125a | hsa-mir-381 | hsa-mir-574 | hsa-mir-126 | hsa-mir-378b | hsa-mir-101 |  |
| hsa-mir-20a | hsa-mir-199a-2 | hsa-mir-125b-2 | hsa-mir-382 | hsa-mir-576 | hsa-mir-184 | hsa-mir-378e | hsa-mir-124 |  |
| hsa-mir-21 | hsa-mir-199b | hsa-mir-126 | hsa-mir-383 | hsa-mir-589 | hsa-mir-206 | hsa-mir-181 | hsa-mir-105 |  |
| hsa-mir-22 | hsa-mir-203a | hsa-mir-127 | hsa-mir-340 | hsa-mir-590 | hsa-mir-375 | hsa-mir-4449 | hsa-mir-7 |  |
| hsa-mir-23a | hsa-mir-204 | hsa-mir-134 | hsa-mir-330 | hsa-mir-615 | hsa-mir-326 | hsa-mir-484 | hsa-mir-103a |  |
| hsa-mir-24-1 | hsa-mir-205 | hsa-mir-136 | hsa-mir-328 | hsa-mir-625 | hsa-miR-324-5p | hsa-mir-1273g | hsa-mir-26a |  |
| hsa-mir-24-2 | hsa-mir-210 | hsa-mir-146a | hsa-mir-342 | hsa-mir-627 | hsa-miR-133b | hsa-mir-675 | hsa-mir-24 |  |
| hsa-mir-25 | hsa-mir-211 | hsa-mir-149 | hsa-mir-337 | hsa-mir-629 | hsa-miR-429 | hsa-mir-1271 | hsa-mir-19b |  |
| hsa-mir-26a-1 | hsa-mir-212 | hsa-mir-150 | hsa-mir-151a | hsa-mir-33b | hsa-miR-449a | hsa-mir-1303 | hsa-mir-196a |  |
| hsa-mir-26b | hsa-mir-181a-1 | hsa-mir-154 | hsa-mir-135b | hsa-mir-652 | hsa-miR-451a | hsa-mir-320e | hsa-mir-135a |  |
| hsa-mir-27a | hsa-mir-214 | hsa-mir-185 | hsa-mir-148b | hsa-mir-449b | hsa-miR-612 | hsa-mir-618 | hsa-mir-181b |  |
| hsa-mir-28 | hsa-mir-215 | hsa-mir-186 | hsa-mir-331 | hsa-mir-411 | hsa-miR-219a-1-3p | hsa-mir-421 | hsa-mir-9 |  |
| hsa-mir-29a | hsa-mir-216a | hsa-mir-188 | hsa-mir-324 | hsa-mir-654 | hsa-mir-130b | hsa-mir-1180 | hsa-let-7f |  |
| hsa-mir-30a | hsa-mir-218-1 | hsa-mir-190a | hsa-mir-338 | hsa-mir-660 | hsa-miR-130b-5p | hsa-mir-613 | hsa-mir-153 |  |
| hsa-mir-31 | hsa-mir-219-1 | hsa-mir-193a | hsa-mir-339 | hsa-mir-542 | hsa-miR-500a-5p | hsa-mir-650 | hsa-mir-667 |  |
| hsa-mir-32 | hsa-mir-219a-1 | hsa-mir-194-1 | hsa-mir-335 | hsa-mir-767 | hsa-mir-665 | hsa-mir-1228 | hsa-mir-103 |  |
| hsa-mir-33a | hsa-mir-221 | hsa-mir-195 | hsa-mir-345 | hsa-mir-1224 | hsa-mir-106 | hsa-mir-1290 | hsa-mir-1205 |  |
| hsa-mir-92a-2 | hsa-mir-222 | hsa-mir-200c | hsa-mir-196b | hsa-mir-1296 | hsa-mir-486-1 | hsa-mir-27 | hsa-mir-44352 |  |
| hsa-mir-93 | hsa-mir-223 | hsa-mir-155 | hsa-mir-423 | hsa-mir-454 | hsa-mir-217 | hsa-mir-577 | hsa-mir-6888 |  |
| hsa-mir-95 | hsa-mir-224 | hsa-mir-181b-2 | hsa-mir-424 | hsa-mir-320b-2 | hsa-mir-1297 | hsa-mir-320c-1 | hsa-mir-4691 |  |
| hsa-mir-96 | hsa-mir-200b | hsa-mir-128b | hsa-mir-425 | hsa-mir-874 | hsa-mir-30 | hsa-mir-543 | hsa-mir-6512 |  |
| hsa-mir-98 | hsa-let-7g | hsa-mir-194-2 | hsa-mir-18b | hsa-mir-708 | hsa-mir-638 | hsa-mir-122a | hsa-mir-1291 |  |
| hsa-mir-99a | hsa-let-7i | hsa-mir-106b | hsa-mir-20b | hsa-mir-744 | hsa-mir-199a | hsa-mir-548f | hsa-mir-3118 |  |
| hsa-mir-100 | hsa-mir-1-2 | hsa-mir-29c | hsa-mir-431 | hsa-mir-873 | hsa-mir-371a | hsa-mir-466 | hsa-mir-4730 |  |
| hsa-mir-101-1 | hsa-mir-15b | hsa-mir-30c-1 | hsa-mir-433 | hsa-mir-1249 | hsa-mir-513c | hsa-mir-350 | hsa-mir-291 |  |
| hsa-mir-29b-1 | hsa-mir-23b | hsa-mir-200a | hsa-mir-409 | hsa-mir-1307 | hsa-mir-571 | hsa-mir-384 | hsa-mir-669c |  |
| hsa-mir-29b-2 | hsa-mir-27b | hsa-mir-302a | hsa-mir-376b | hsa-mir-513b | hsa-mir-1246 | hsa-mir-3 | hsa-mir-4 |  |
| hsa-mir-103-2 | hsa-mir-30b | hsa-mir-101-2 | hsa-mir-483 | hsa-let-7a-5p | hsa-mir-1825 | hsa-mir-602 | hsa-mir-85 |  |
| hsa-mir-103a-2 | hsa-mir-122 | hsa-mir-34b | hsa-mir-485 | hsa-let-7f-5p | hsa-mir-544a | hsa-mir-520b | hsa-mir-5703 |  |
| hsa-mir-103-1 | hsa-mir-124-1 | hsa-mir-34c | hsa-mir-488 | hsa-miR-16-5p | hsa-mir-885 | hsa-mir-9500 | hsa-mir-5120 |  |
| hsa-mir-103a-1 | hsa-mir-124-2 | hsa-mir-299 | hsa-mir-490 | hsa-miR-17-5p | hsa-mir-1207 | hsa-mir-1182 | hsa-mir-6881 |  |
| hsa-mir-106a | hsa-mir-124-3 | hsa-mir-301a | hsa-mir-491 | hsa-mir-17* | hsa-mir-1301 | hsa-mir-4284 | hsa-mir-7240 |  |
| hsa-mir-16-2 | hsa-mir-125b-1 | hsa-mir-99b | hsa-mir-146b | hsa-miR-17-3p | hsa-mir-29 | hsa-mir-875 | hsa-mir-1261 |  |
| hsa-mir-192 | hsa-mir-128-1 | hsa-mir-296 | hsa-mir-202 | hsa-miR-19b-3p | hsa-mir-4739 | hsa-mir-3666 | hsa-mir-295 |  |
| hsa-mir-197 | hsa-mir-130a | hsa-mir-130b | hsa-mir-493 | hsa-miR-24-3p | hsa-let-7 | hsa-mir-513a | hsa-mir-4481 |  |
| hsa-mir-199a-1 | hsa-mir-132 | hsa-mir-30e | hsa-mir-432 | hsa-miR-26a-5p | hsa-mir-148 | hsa-mir-1268a | hsa-mir-1968 |  |
| hsa-mir-208a | hsa-mir-133a-1 | hsa-mir-26a-2 | hsa-mir-494 | hsa-miR-32-5p | hsa-mir-216b | hsa-mir-619 | hsa-mir-4791 |  |

**Figure 1.**
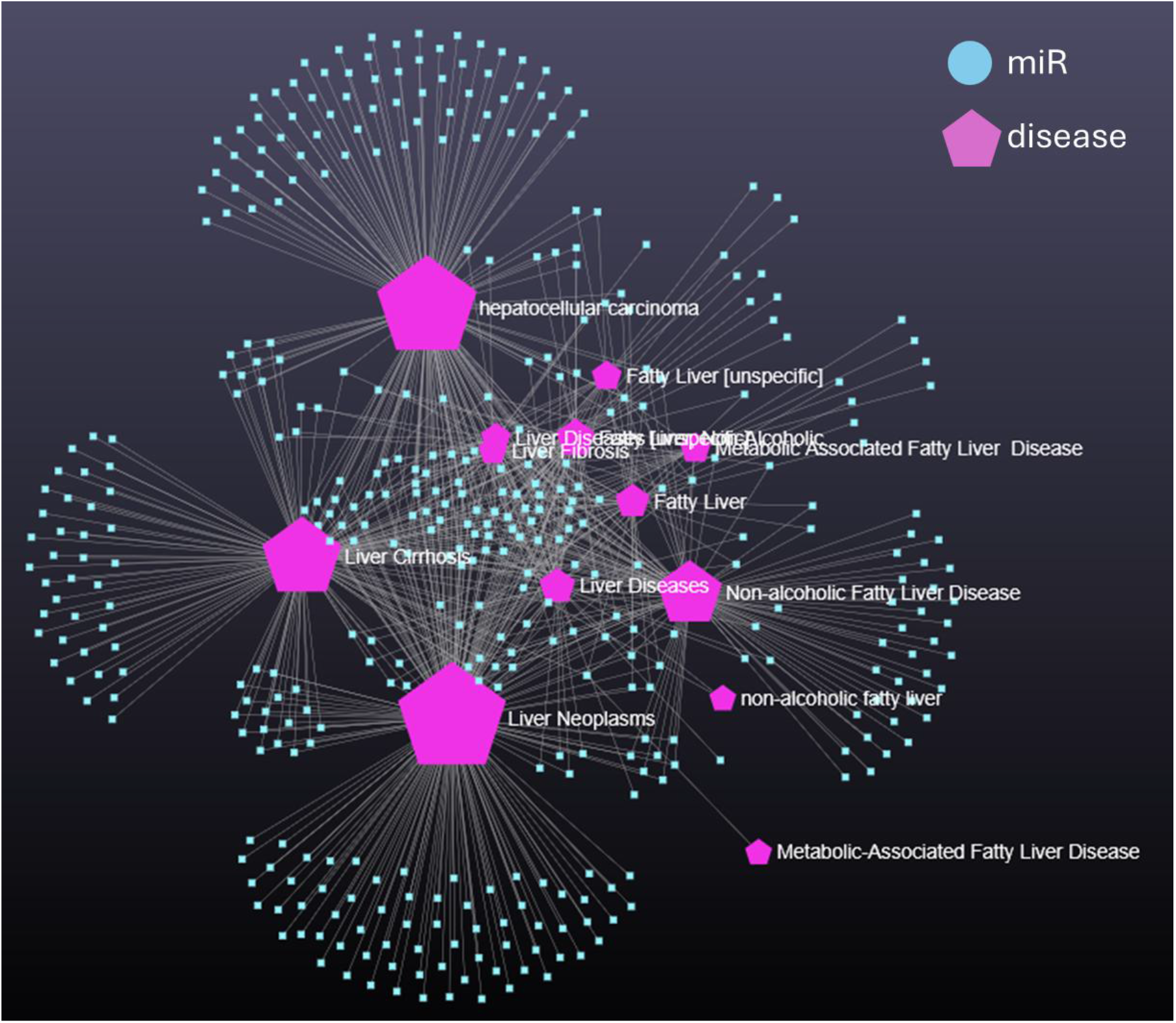
The MASLD-HCC Interactome. Forced atlas representations of the miRs associated with MASLD-related HCC. More than 400 miRs comprise this interactome (Table 1).

Next, we identified miRs regulating biomarkers that support a diagnosis of HCC. genes for AFP, prothrombin/DCP/PIVKA II, glypican-3, OPN, GP73 and DKK1 are regulated by 274 miRs (Table 2).

**Table 2.**
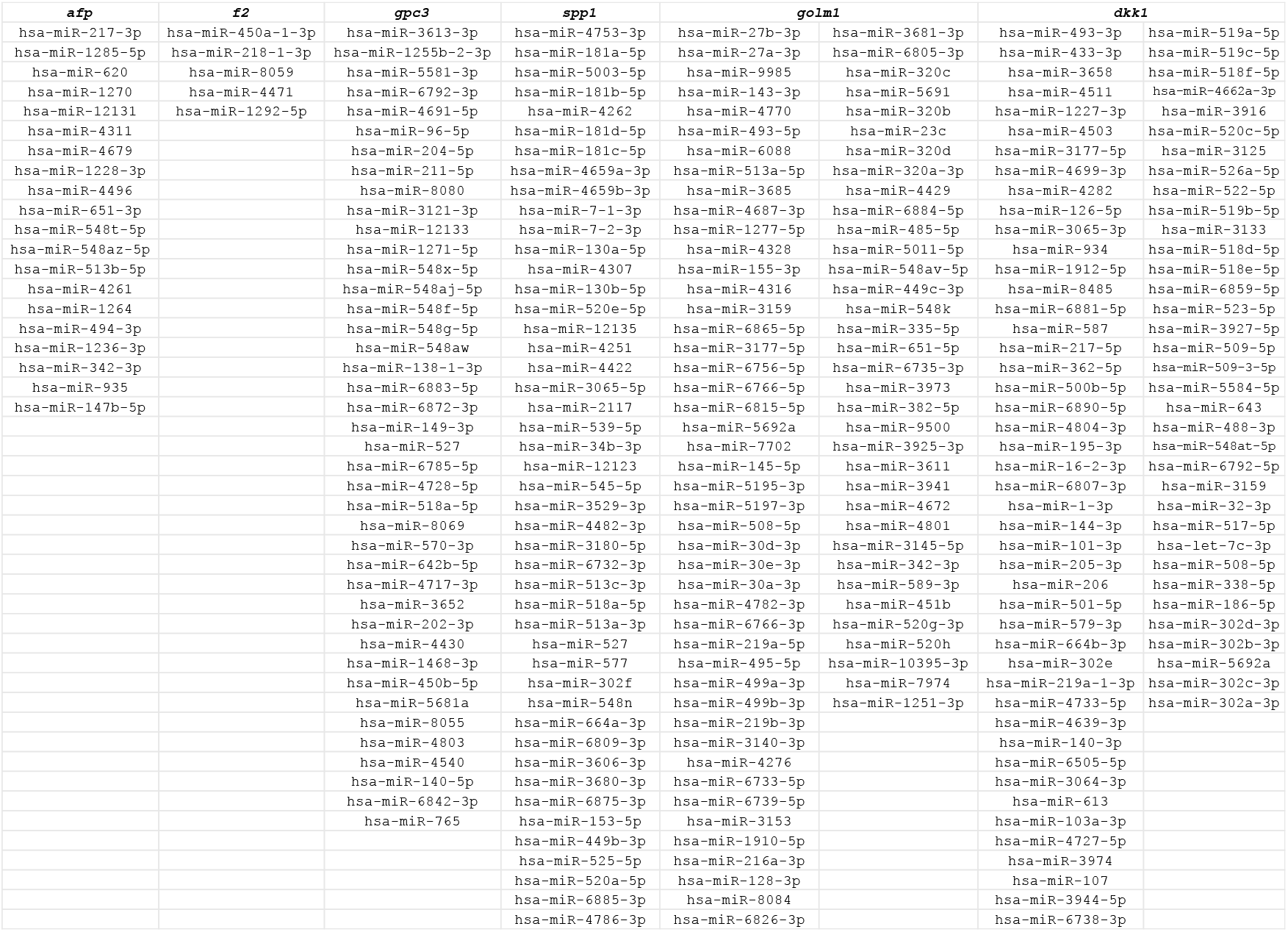
Library of miRs Regulating Genes Associated with HCC Biomarkers.

A total of 9 miRs were common to both libraries. These were hsa-miR-577, hsa-miR-130b-5p, hsa-miR-9500, hsa-miR-101-3p, hsa-miR-206, hsa-miR-219a-13p, hsa-miR-613, hsa-miR-103a- 3p and hsa-miR-107. Of these, literature indicates that hsa-miR-577, hsa-miR-9500, hsa-miR-101- 3p, has-miR-206, hsa-miR-219a-13p and hsa-miR-613 exhibit tumor suppressive activity with their downstream gene products exhibit oncogenic potential.

miR-577 is frequently downregulated in colorectal and breast cancers, where it suppresses tumor cell proliferation, invasion, and EMT^21^. Mechanistically, miR-577 targets oncogenic factors such as HSP27 in colorectal cancer and RAB25 in breast cancer, leading to reduced metastatic potential^22^. OPN, regulated by this miR, interacts with integrins including αvβ3 and CD44, stimulating PI3K/Akt, NF-κB, and MAPK pathways, which promote cell motility, invasiveness, and survival^23^. In HCC, OPN is overexpressed^5^, and it can drive a cancer stem cell-like phenotype through the integrin–NF-κB–HIF-1α axis, enhancing self-renewal and therapy resistance. It recruits and alters tumor-associated macrophages and fibroblasts, aiding immune evasion and angiogenesis^24^. OPN is broadly overexpressed in many solid tumors where similar signaling promotes metastatic spread, angiogenesis, and immunosuppression^25^. Elevated levels of OPN in plasma often correlate with poor prognosis and advanced disease in HCC^5^.

miR-9500 is significantly downregulated in lung cancer, where its restoration suppresses proliferation, migration, and invasion^26^. Functionally, miR-9500 directly targets AKT1, resulting in inhibition of PI3K/AKT signaling and reduced tumor growth *in vitro* and *in vivo*^*26*^. GP73 regulated by hsa-miR-9500, resides in the Golgi and influences processing and secretion of proteins, including cancer-relevant growth factors and enzymes^27^. Its overexpression is linked to enhanced proliferation, invasion, metastasis, and may facilitate secretion of factors that strengthen tumor survival and microvascular growth^28^. Some evidence suggests GP73 affects immune signals and angiogenesis with GP73 levels elevated in several cancers including HCC, prostate and lung. GP73 is overexpressed in HCC and is used as a biomarker for HCC and investigated for detection of other cancers^5,27,28^; it’s also considered as a therapeutic target for blocking pro-tumor secretory pathways^5^.

miR-101-3p is a well-established tumor suppressor in HCC and is consistently downregulated in tumor tissue^29^. It’s overexpression inhibits HCC cell proliferation, migration, invasion, and EMT by targeting HGF/c-Met, BIRC5 (survivin), and downstream PI3K–AKT signaling^30^. Its tumor-suppressive function is also conserved across multiple cancer types, supporting a central role in oncogenic pathway regulation^31^. miR-206 is frequently downregulated in HCC and acts as a tumor suppressor by inhibiting proliferation, migration, and tumor progression^32,33^. It targets oncogenic drivers including c-Met and regulators of the cell cycle, leading to suppression of growth and metastatic phenotypes^33^. Similar tumor-suppressive effects are observed in lung and other cancers, highlighting a conserved anti-oncogenic function^32,33^. miR- 219a-1-3p is downregulated in colon and gastric cancers, where it inhibits proliferation, migration, and invasion^34^. Mechanistically, it targets MEMO1, a regulator of cytoskeletal dynamics and metastatic signaling3^4,35^. While direct evidence in HCC is currently absent, its role in suppressing EMT-related phenotypes suggests possible relevance in liver cancer progression^34,35^. miR-613 is broadly recognized as a tumor suppressor and is downregulated in several malignancies, including HCC^36,37^. In liver cancer contexts, miR-613 suppresses tumorigenesis by targeting oncogenic drivers such as DCLK1, reducing proliferation and invasive capacity^38^. Similar tumor-suppressive effects are reported in lung, colorectal, and other cancers, where miR-613 enhances chemosensitivity and limits metastasis^39^. miR-101-3p, miR-206 and miR-613 regulate expression of *dkk1*, the DKK1 gene. Elevated DKK1 levels have been reported in HCC^5^; DKK1 promotes hepatocellular carcinoma cell migration and invasion through β-catenin/MMP7 signaling pathway^40^ and its expression levels correlate with worse prognosis in several cancers and are under study as both biomarker and therapeutic target.

## Discussion

In the present study we identify hsa-miR-577, hsa-miR-9500, hsa-miR-101-3p, miR-206, hsa-miR-219a-1-3p, and hsa-miR-613 with tumor suppressive properties and potentially reduced expression levels in HCC. OPN, GP73 and DKK1, gene products under regulation by these miRs, carry oncogenic potential and exhibit increased expression levels in HCC. Together, these findings support a model in which reduced tumor-suppressive miR control contributes to oncogenic biomarker overexpression and progression from MASLD to HCC.

The central observation that drove this study is that the liver uniquely converts chronic metabolic injury and fibrosis into malignant transformation at a far higher frequency than other organs with end-stage fibrotic disease^42^. Our findings suggest that this disparity may, at least in part, reflect liver-specific dysregulation of miR–mRNA networks, that simultaneously amplify oncogenic signaling. The liver’s unique metabolic, immunological, and regenerative milieu may magnify the consequences of miR loss, tipping fibrotic repair toward neoplastic transformation.

OPN, GP73, and DKK1 emerge as particularly relevant nodes within this regulatory network. OPN promotes EMT, angiogenesis, immune cell recruitment, and cancer stem cell–like properties, all of which are hallmarks of aggressive disease and therapeutic resistance. The regulation of OPN by tumor-suppressive hsa-miR-577 suggests that miR downregulation in MASLD may enable sustained OPN overexpression, thereby reinforcing a pro-tumorigenic microenvironment as fibrosis advances. GP73, a Golgi-resident protein widely used as a circulating HCC biomarker, represents a second mechanistic axis linking miR dysregulation to malignant progression. Although GP73 is classically viewed as a diagnostic marker, growing evidence indicates that it actively participates in tumor biology by modulating protein trafficking, secretion of growth factors, and possibly immune signaling. A reduction in expression levels of tumor-suppressive hsa-miR-9500–mediated repression may facilitate aberrant GP73 expression, enhancing secretory pathways that support tumor growth, angiogenesis, and invasion. In MASLD, where hepatocytes experience sustained metabolic and endoplasmic reticulum stress, dysregulated Golgi function may further amplify GP73-driven oncogenic effects. DKK1 only reinforces this program. Its regulation by tumor-suppressive miRs, hsa-miR-101-3p, hsa-mir-206, hsa-miR-219a- 13p and miR-613 highlights how miR expression loss may simultaneously deregulate developmental pathways, extracellular matrix remodeling, and immune surveillance in the cirrhotic liver. Indeed, inclusion of the miRs in the MASLD-HCC interactome reinforces the concept that metabolic liver disease converges on canonical oncogenic pathways via miR dysregulation.

From a translational perspective, these findings have several implications. First, coordinated measurement of tumor-suppressive miRs alongside established circulating biomarkers such as OPN, GP73, and DKK1 may improve early detection and risk stratification of HCC in MASLD patients, particularly in those without overt cirrhosis. Second, restoration of tumor-suppressive miRs and/or targeting of their oncogenic gene products, represent a potential therapeutic strategy to interrupt progression along the MASLD continuum. Finally, these miR– biomarker axes may help explain why a subset of MASLD patients develop HCC in the absence of advanced fibrosis, underscoring the need for molecular, rather than purely histologic, risk assessment.

In summary, our integrative analysis identifies a miR–gene regulatory network that links metabolic liver disease to oncogenic biomarker expression and malignant transformation. Dysregulation of tumor-suppressive miRs and consequent overexpression of OPN, GP73, and DKK1 may act as key drivers of the MASLD-to-HCC transition. Further mechanistic studies and longitudinal clinical validation will be essential to determine whether these miRs and their targets can serve as predictive biomarkers or therapeutic entry points in this rapidly expanding patient population.

## Conclusion

Mechanistic insights should illuminate therapies. The nodes and their pathways identified herein not only illuminate a potential mechanistic basis for the transition of MASLD to HCC but also inform an array of targets to mitigate this transition.

## Notes

### Competing Interest Statement

The authors have declared no competing interest.

## References

1) Miao L et al. Current status and future trends of the global burden of MASLD. Trends in Endocrinology and Metabolism, 2024, 35:697–707.

2) Rinella ME et al. A multisociety Delphi consensus statement on new fatty liver disease nomenclature. Journal of Hepatology, 2023, 79:1542–1556.

3) Obesity and overweight; last accessed, 12/15/24.

4) Diabetes; last accessed, 12/15/24.

5) Hwang A et al. Supervised learning reveals circulating biomarker levels diagnostic of hepatocellular carcinoma in a clinically relevant model of non-alcoholic steatohepatitis; An OAD to NASH. 2018, 10.1371/journal.pone.0198937

6) HCC in metabolic dysfunction-associated steatotic liver disease | AASLD; last accessed 12/23/24.

7) Tarao K et al. Real impact of liver cirrhosis on the development of hepatocellular carcinoma in various liver diseases—meta-analytic assessment. Cancer Med 2019, 8:1054–1065.

8) Mittal S et al. Hepatocellular Carcinoma in the Absence of Cirrhosis in United States Veterans is Associated With Nonalcoholic Fatty Liver Disease. Clin Gastroenterol Hepatol 2016, 14:124-31.e1

9) Kanwal F et al. Risk of Hepatocellular Cancer in Patients with Non-alcoholic Fatty Liver Disease. Gastroenterology 2018, 155:1828–1837.e2

10) Piscaglia, F et al.Clinical Patterns of Hepatocellular Carcinoma in Nonalcoholic Fatty Liver Disease: A Multicenter Prospective Study. Hepatology 2016, 63:827–838.

11) Stine JG et al. Systematic review with meta-analysis: risk of hepatocellular carcinoma in non-alcoholic steatohepatitis without cirrhosis compared to other liver diseases, Aliment Pharmacol Ther Aliment Pharmacol Ther 2018, 48:696–703.

12) Ertle J et al. Non-alcoholic fatty liver disease progresses to hepatocellular carcinoma in the absence of apparent cirrhosis. Int J Cancer 2011, 128:2436–43.

13) Vaz J et al. Metabolic dysfunction-associated steatotic liver disease has become the most common cause of hepatocellular carcinoma in Sweden: A nationwide cohort study. Int J Cancer 2025, 156:40–51.

14) Bastos VAF et al. Shared and Context-Specific Mechanisms of EMT and Cellular Plasticity in Cancer and Fibrotic Diseases. Int J Mol Sci 2025, 26:9476

15) Maher TM. Interstitial Lung Disease: A Review. JAMA. 2024, 331:1655–1665

16) Deng L. et al.Global, regional, and national burden of chronic kidney disease and its underlying etiologies from 1990 to 2021: a systematic analysis for the Global Burden of Disease Study 2021. BMC Public Health 2025, 25:636.

17) Cojan-Minzat BO et al. Non-ischemic dilated cardiomyopathy and cardiac fibrosis. Heart Fail Rev 2021, 26:1081–1101.

18) Chery G et al. Prognostic value of myocardial fibrosis on cardiac magnetic resonance imaging in patients with ischemic cardiomyopathy: A systematic review. Am Heart J. 2020, 229:52–60.

19) Al-Hasan M et al. Role of biomarkers in the diagnosis and management of HCC. Liver Transpl 2025, 31:384–394.

20) Bissoondial TL et al. Identification of disease-associated microRNA in a diet-induced model of nonalcoholic steatohepatitis. Mol Omics 2021,17:911-916.

21) Jiang H et al. microRNA-577 suppresses tumor growth and enhances chemosensitivity in colorectal cancer. J Biochem Mol Toxicol 2017, 31.

22) Yin C et al. MiR-577 suppresses epithelial-mesenchymal transition and metastasis of breast cancer by targeting Rab25. Thorac Cancer 2018, 9:472–479.

23) Kundu JK and Surh- YJ. Emerging avenues linking inflammation and cancer. Free Radical Biology and Medicine 2012, 52:2013–2037.

24) Bastos ACSDF et al. The Intracellular and Secreted Sides of Osteopontin and Their Putative Physiopathological Roles. J Mol Sci 2023, 24:2942.

25) Kundu G and Elangovan S. Investigating the Role of Osteopontin (OPN) in the Progression of Breast, Prostate, Renal and Skin Cancers. Biomedicines 2025, 13:173.

26) Yoo JK et al. The novel miR-9500 regulates the proliferation and migration of human lung cancer cells by targeting Akt1. Cell Death & Differentiation 2014, 21:1150–1159.

27) Wang Y and Wan YJY. Golgi protein 73, hepatocellular carcinoma and other types of cancers. Liver Res 2020, 4:161–167.

28) Liu Y et al. GP73-mediated secretion of AFP and GP73 promotes proliferation and metastasis of hepatocellular carcinoma cells. Oncogenesis 2021, 10:69.

29) Liu Z et al. MicroRNA-101 suppresses migration and invasion via targeting vascular endothelial growth factor-C in hepatocellular carcinoma cells. Oncol Lett 2015, 11:433– 438.

30) Zhu W et al. MiR-101-3p targets the PI3K-AKT signaling pathway via Birc5 to inhibit invasion, proliferation, and epithelial–mesenchymal transition in hepatocellular carcinoma. Clin Exp Med 2025, 25:88.

31) Liu P et al. mir-101-3p is a key regulator of tumor metabolism in triple negative breast cancer targeting AMPK. Oncotarget 2016, 7:35188–35198.

32) MicroRNA-206 as a promising epigenetic approach to modulate tumor-associated macrophages in hepatocellular carcinoma World J Gastroenterol 2024, 30:4503–4508.

33) MicroRNA-206 inhibits the viability and migration of human lung adenocarcinoma cells partly by targeting MET. Oncol Lett 2016, 12:1171–117.

34) Luo M et al. MiRNA-219a-1-3p inhibits the malignant progression of gastric cancer and is regulated by DNA methylation. Oncologie 2023, 25: 495–506.

35) Xu K et al. miR-219a-1 inhibits colon cancer cells proliferation and invasion by targeting MEMO1. Cancer Biol Ther 2020, 21:1163–1170.

36) Li B et al. miR-613 inhibits liver cancer stem cell expansion by regulating SOX9 pathway. Gene 2019, 707:78–85.

37) Jiang X et al. MiR-613 functions as tumor suppressor in hepatocellular carcinoma by targeting YWHAZ. Gene 2018, 659:168–174.

38) Wang W et al. miR-613 inhibits the growth and invasiveness of human hepatocellular carcinoma via targeting DCLK1. Biochem Biophys Res Commun 2016, 473:987–992.

39) Mei J et al. MicroRNA-613: A novel tumor suppressor in human cancers. Biomedicine & Pharmacotherapy 2020, 123:109799.

40) Chen L et al. DKK1 promotes hepatocellular carcinoma cell migration and invasion through β-catenin/MMP7 signaling pathway. Mol Cancer 2013, 12:157.

41) Wei R et al. Analyzing the prognostic value of DKK1 expression in human cancers based on bioinformatics. Ann Transl Med 2020, 8:552.

42) Affo S et al. The Role of Cancer-Associated Fibroblasts and Fibrosis in Liver Cancer. Annu Rev Pathol. 2016, 12:153–186.

